# Targeting lateral inhibition to improve vision following macular degeneration

**DOI:** 10.1101/2020.02.21.953828

**Authors:** M Rizzi, K Powell, MR Robinson, T Matsuki, J Hoke, RN Maswood, A Georgiadis, M Georgiou, PR Jones, C Ripamonti, M Michaelides, GS Rubin, AJ Smith, RR Ali

**Affiliations:** UCL Institute of Ophthalmology, London, UK; CRS Ltd. Rochester, UK; NIHR Biomedical Research Centre at Moorfields Eye Hospital and UCL Institute of Ophthalmology

## Abstract

Macular degeneration is the leading cause of blindness in the developed world. Whilst most patients lose sight owing to atrophic changes, no treatments currently exist that improve the vision deficit due to atrophy. Here, we identify loss of lateral inhibition as a specific mechanism by which photoreceptor degeneration reduces visual function beyond the atrophic area. We find that this inhibition is adaptive, and that if modulated can improve visual function, making inhibitory circuits an unexpected therapeutic target for age related macular degeneration and related disorders.

## Main text

Visual circuits comprise lateral as well as feed-forward connections^1^. Altering the amount of lateral input greatly changes contrast sensitivity and spatial acuity, highlighting the importance of lateral connections in visual function (Fig.1a,b,c; Suppl.Fig1-3). We demonstrate for the first time that in normal vision, surround luminance acts as an adaptive mechanism that optimizes vision to contextual luminance (Fig.1d; Suppl.Fig.4). We therefore reasoned that local photoreceptor degeneration would result in adjacent areas being deprived of an adaptive input and thus manipulating laterally projecting inhibitory circuits may improve vision following photoreceptor loss (Suppl.Fig.5).

**Figure 1.**
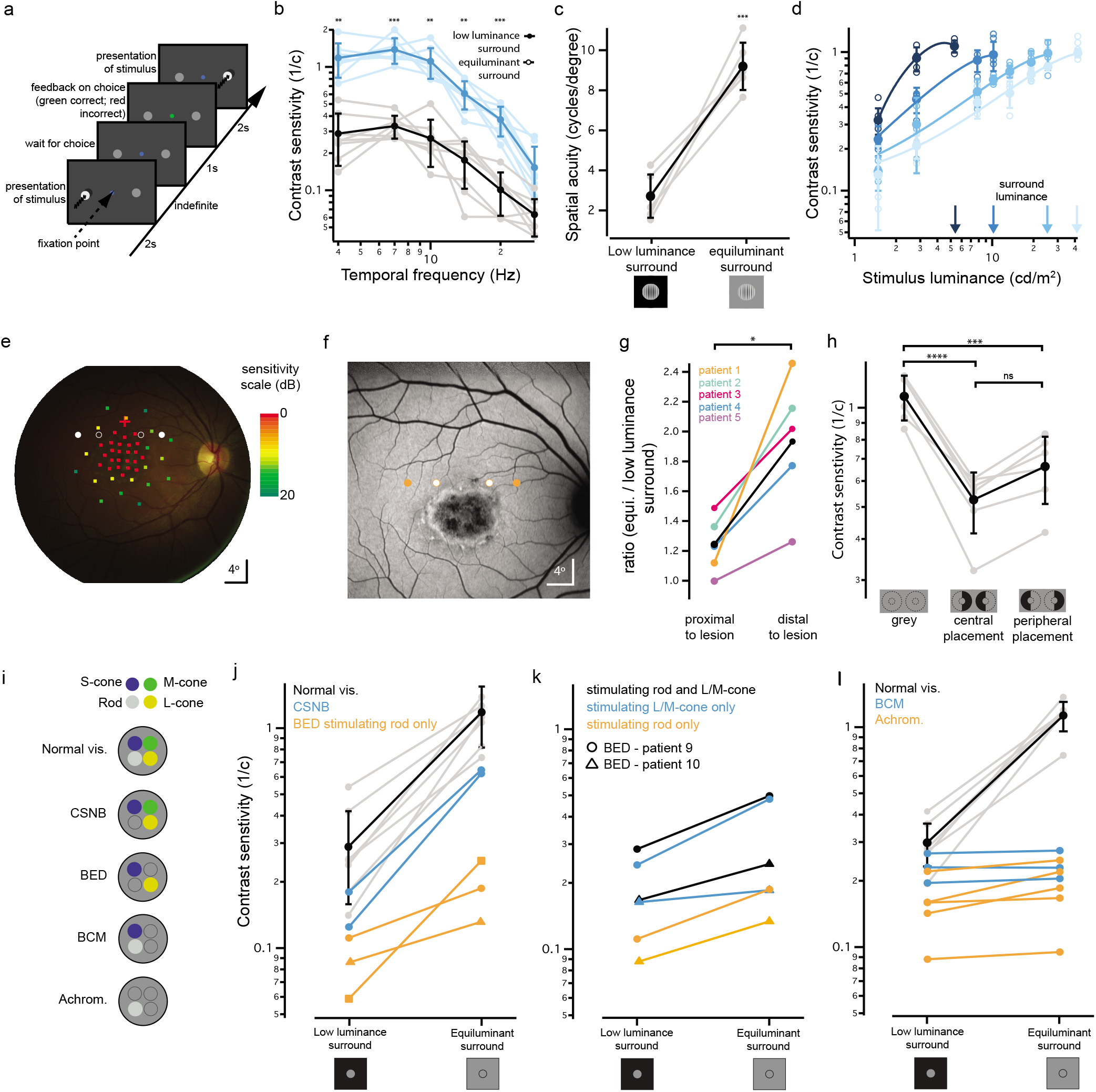
Equiluminant surround input improves contrast encoding in human subjects. **a**, Schematic of the experimental procedure. A 2-alternative forced choice task was used to measure contrast sensitivity thresholds. Stimuli were either a temporally or spatially modulated stimulus presented on either side of the fixation spot. Subjects were asked to indicate on which side the stimulus was presented. Their ability to detect the stimulus was determined by their forced choice response. **b,c**, Vision was improved by an equiluminant (27.7cd/m^2^) vs low luminance (0.15cd/m^2^) surround in normal vision subjects. Data for temporal contrast sensitivity shown in b (n=8). Data for visual acuity shown in c (n=6). The difference is equivalent to a 0.52-to-1.03 shift on the logMAR scale (n=6). **d**, The effect of several surround luminance values (blue arrows) were compared for a range of mean stimulus luminance values. Contrast encoding functions (corresponding blue color) were shifted rightwards with increasing surround luminance, but maximum contrast sensitivity remained comparable (n=4). **e**, Microperimetry allowed functional mapping of the degenerating area in patients with Stargardt disease. Color encoded microperimetry scores (in dB) are shown here superimposed on a fundus reflectance image. Red cross indicates average fixation location. Scores guided the placement of stimuli to allow comparison of contrast sensitivity at a location proximal to the lesion (open circles) with a location more distal to the lesion (filled circles) for each patient. **f**, Fundus Autofluorescence imaging (486 nm) of the retina presented in panel d, showing the extent of the macular lesion. Open and filled circles indicate the proximal and distal locations tested. **g**, Comparison of contrast thresholds in five patients with Stargardt disease, each indicated with a different color. Measurements were made at locations proximal and distal to the lesion, and when surrounded with low or equiluminant surrounds. Data is expressed as a ratio of contrast sensitivity for equiluminant over low luminance surrounds at each location for each patient. **h**, An artificial scotoma created by placing a low luminance half annulus adjacent to the test stimulus significantly worsened contrast sensitivity thresholds in normal vision subjects (n=6). **i**, Schematic showing the contributing photoreceptor types in normal vision subjects, congenital stationary night blindness (CSNB, cone-only vision), Bornholm eye disease (BED, lack of one cone opsin; M-opsin for two of three patients tested, L-opsin for one patient) blue cone monochromatism (BCM, rod and S-cone vision), achromatopsia (rod-only vision). **j**, Selective stimulation of rod photoreceptors in BED patients (orange traces) or cone photoreceptors (CSNB patients, blue traces) showed a positive rather than suppressive effect. Comparative data for normal vision subjects is shown in black (n=8, data from Fig.1b). **k**, An equiluminant surround improved vision in two Bornholm eye disease patients using rod isolating (orange traces), cone isolating (blue traces) and rod-cone mediated vision (black traces). **l**, An equiluminant surround provided only a marginal improvement over a mismatched surround in patients with achromatopsia (n=5, orange traces) and blue cone monochromatism (n=3, blue traces), indicating an essential role for L/M cone photoreceptors in mediating the effect. All contrast sensitivity values are calculated as 1/contrast threshold (1/c). Error bars denote standard deviations around a mean values. *p*-values were calculated using a one-way ANOVA, with Tukey’s post hoc multiple comparison test (b); Dunnet’s post-hoc multiple comparisons test (h); or a paired t-test (c, g). \*\*\*\**p*<0.0001 \*\*\**p*<0.001, \*\**p*<0.01, \**p*<0.05.

To assess the impact on vision in macular degeneration, we recruited a group of subjects with Stargardt disease^2^. Microperimetry testing provided functional mapping of the central retina (Fig.1e; Suppl.Fig.6). This guided the positioning of stimuli to test contrast sensitivity in locations adjacent to and further away from the atrophic area for each patient (Fig. 1f; Suppl.Fig.6). Not surprisingly, contrast sensitivity in Stargardt patients was poor compared to normal vision subjects and it was poorer at locations close to the lesion compared with further away from it. To measure the effect of the lateral adaptive input, we performed a specific comparison between contrast sensitivity thresholds within each location when surrounded by either an equiluminant or a low luminance field. For stimuli presented distal to degenerating areas, patients showed a surround-mediated gain in contrast sensitivity that was significantly larger than for adjacent locations (Fig. 1g, Suppl.Fig.7). We confirmed this finding in normal vision subjects, by using an artificial scotoma to cause a reduction in contrast sensitivity (Fig.1h). These results show that a scotoma affects visual processing beyond its border, significantly worsening the quality of remaining vision. Importantly, the effect of lateral feedback on quality of vision is substantial (Fig.1b,c,g,h,j), equivalent, in terms of visual acuity, to five lines on a logMAR chart.

To find candidate therapeutic targets we probed the cell types that underlie these lateral interactions. We recruited subjects with rare genetic conditions to differentially assess the role of L, M and S-cone, and rod photoreceptors in lateral adaptation (Fig.1i). Lateral input improved contrast sensitivity in subjects with cone-only vision (congenital stationary night blindness^3^, Fig.1j, blue line; Suppl.Fig.8) and in patients where single cone classes could be stimulated in isolation using silent substitution^4^ (Bornholm eye disease^5^, Fig.1k, blue lines). Interestingly, we found a similar effect when using rod-isolating stimuli (Fig.1j,k orange lines; Suppl.Fig.9), revealing that lateral adaptation optimizes both cone and rod mediated vision. In patients with achromatopsia and with blue cone monochromatism, who lack function in all L/M-cones^5^, the effect of the lateral input gain mechanism was absent (Fig. 1l), determining that L/M-cones are the source of lateral adaptation. These results explain for the first time why a local or global loss of cone function causes light aversion^6^, as cones are unable to adapt rods to mesopic luminance levels. Importantly, we identify cone-driven lateral inhibition as a potential therapeutic target to improve vision.

To directly investigate this, we used a mouse model in which lateral adaptation should be missing. *Cnga3^−/−^* mice have an analogous lack of functioning of cone-associated CNG channels as the achromat patients (Fig. 1l, orange lines). These mice should similarly lack lateral adaptation. To confirm this, we performed two contrast sensitivity tasks with wild-type and *Cnga3^−/−^* mice. We found reduced sensitivity (Fig.2a,b; Suppl.Fig.10) and adaptation to lateral luminance (Fig.2a) in *Cnga3^−/−^* mice, consistent with data in human subjects.

**Figure 2.**
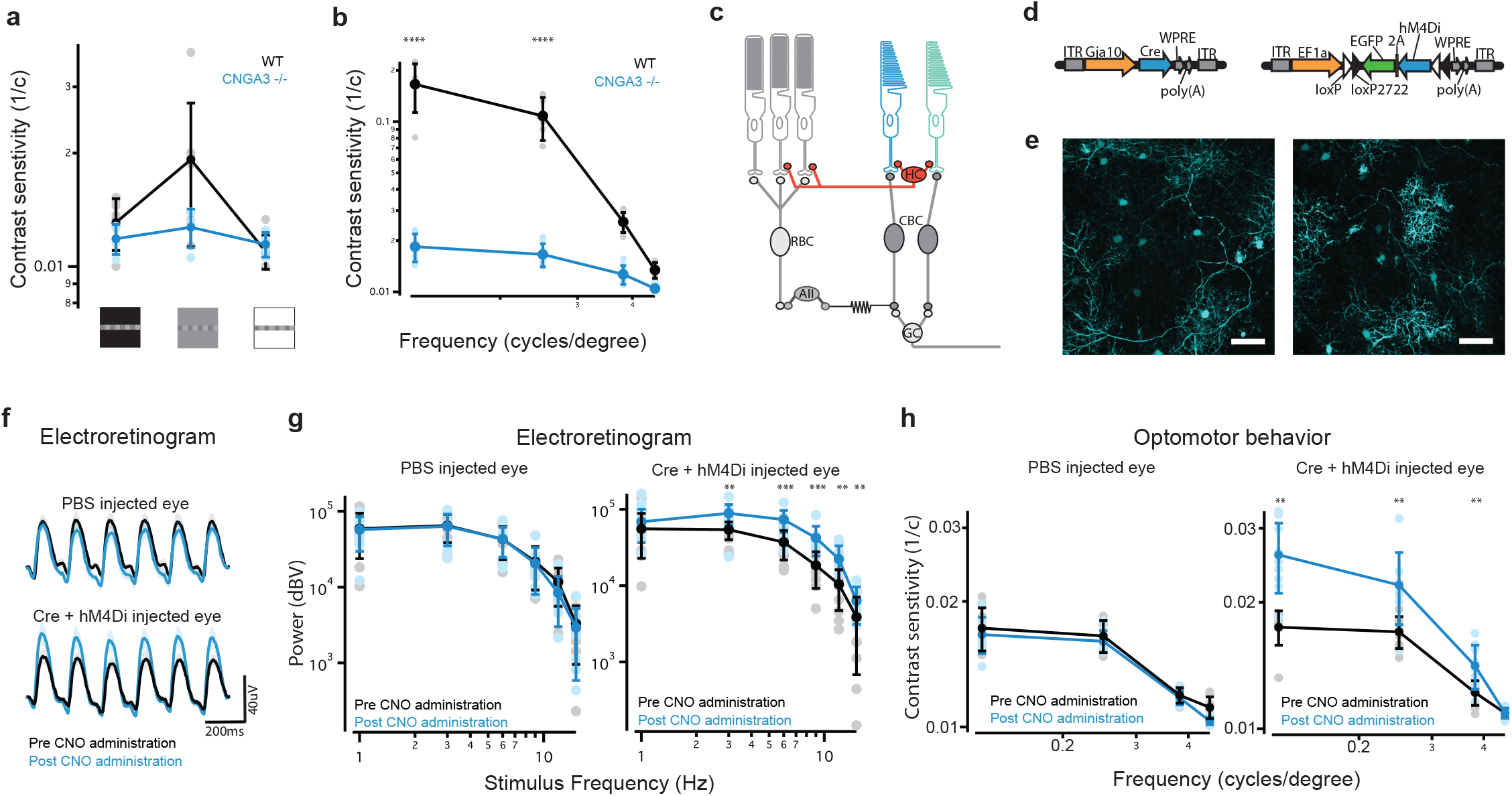
Improved vision following restoration of lateral input to rod photoreceptors. **a**, Surround luminance affects contrast encoding in mice in an optomotor behavior (see Methods). Wild-type mice (black trace, n=9) showed increased contrast sensitivity with an equiluminant surround compared to low- and high-luminance conditions, whereas *Cnga3^−/−^* mice (blue trace, n=4) did not. **b**, Contrast sensitivity was also reduced in *Cnga3^−/−^* mice (n=9) compared with wild-type mice (n=5) in a standard optomotor behavior. **c**, Schematic of mouse retinal circuits. Horizontal cell (HC) feedback circuit is shown in red. S- and M-cones are shown in blue and green respectively. Rod photoreceptors shown in grey. CBC: cone bipolar cell, RBC: rod bipolar cell, AII: AII amacrine cell, GC: ganglion cell. **d**, Expression of hM4Di in horizontal cells was achieved by co-injection of a Gja10-Cre-expressing vector and a floxed hM4Di-GFP-expressing vector. **e**, Example images of hM4Di-GFP expression in horizontal cells in the mouse retina (wholemount, scale bar = 50μm). **f**, Chemogenetic activation of horizontal cells improves outer retina function. Averaged electroretinogram traces recorded during presentation of a 1s stimulus with a 6hz sinusoid modulation. Tests were repeated before (black traces) and after (blue traces) intraperitoneal injection of the hM4Di activator CNO (n=9; mean average, shaded regions denote standard error). **g**, Fourier analysis (see Methods) of electroretinogram responses for stimuli of different temporal frequencies at 0.1 cd/m^2^, before (black traces) and after (blue traces) CNO injection (n=9). **h**, Optomotor behaviour reveals improved contrast sensitivity following hM4Di activation (n=9). Error bars denote standard deviations around mean values. *p*-values were calculated using a one-way ANOVA, with Tukey’s post-hoc multiple comparisons test (a, b); or with a two-way ANOVA, with Sidak’s post-hoc multiple comparisons test (g, h). \*\*\*\**p*<0.0001 \*\*\**p*<0.001, \*\**p*<0.01, \**p*<0.05.

Neurons in the inner retina survive cone and/or rod dystrophy, including in *Cnga3^−/−^* mice. We therefore took an interventional approach in these mice, by targeting an inner retinal neuron to replace missing lateral adaptation and improve vision. One subtype of horizontal cells (H1) is known to mediate interactions between cones and rods in several species, including humans and mice^7–13^ (Fig.2c). H1 cells receive inputs from cones and are thus not activated in *Cnga3^−/−^* mice^8^. We reasoned that restoring horizontal cell activation might improve function, mimicking lateral feedback. To achieve this, we identified a ~3kb region within the *Gja10* promoter^8,14^ and expressed the chemogenetic hyperpolarizing actuator hM4Di in horizontal cells, using an AAV2/8 vector system (Fig.2d,e; average transduction 29% ± 11%). We found that chemogenetic hyperpolarization of H1 cells improved vision both in terms of voltage modulation in photoreceptors and bipolar cells (Fig.2f,g) and behavioral contrast sensitivity measurement (Fig.2h). These results show that the impairment of vision due to lack of lateral adaptive feedback can be partially reversed, by mimicking light input to laterally projecting inhibitory neurons.

In the human retina, we have shown that lateral inhibition adapts both rod and cone photoreceptors (Fig. 1i-l) meaning this strategy could improve perimacular and foveal vision, that is often spared until the late stages of macular degeneration^15^. Restoration of lateral input may also slow the expansion of the degenerating area, since lack of horizontal cell feedback leads to degeneration of rod photoreceptors^16^ and a correspondingly reduced scotopic response^17–19^. The diverse class of laterally projecting inhibitory neurons^1,20^ should be explored further for additional therapeutic targets.

In summary, our findings advance the understanding of the pathology of macular degeneration and identify a novel therapeutic strategy to improve vision.

