## Supplementary Material for "Targeting lateral inhibition to improve vision following macular degeneration"

### Supplementary figure 1

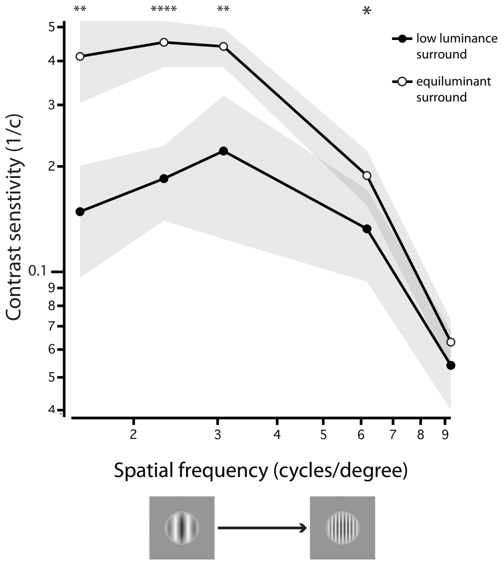


**Surround luminance affects spatial contrast sensitivity functions in normal vision subjects.**
An equiluminant surround improves spatial contrast sensitivity functions. Gabor patches of different spatial frequencies were presented inside a 1° stimulus area, surrounded by a 1.75° annulus width (n=9). All contrast sensitivity values are calculated as 1/contrast threshold (1/c). Values shown are mean averages and shaded regions denote standard deviations. *p*-values were calculated using a one-way ANOVA, with Tukey's post hoc multiple comparison test. *****p*<0.0001, ****p*<0.001, ***p*<0.01, **p*<0.05.

### Supplementary figure 2

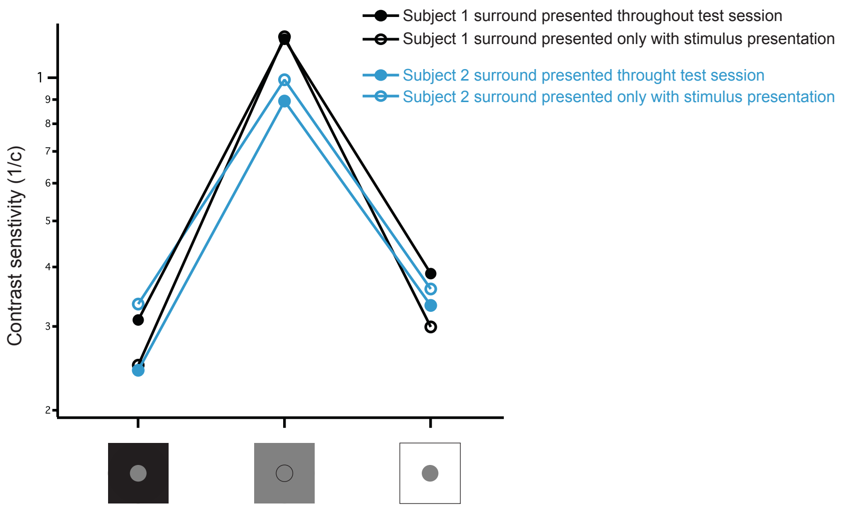


**Surround-mediated effects do not require prolonged adaptation to surround luminance.**
Comparison of temporal contrast sensitivity with low luminance, equiluminant and high luminance surrounds in two normal vision subjects, each indicated with a different color. Contrast thresholds were measured using a 1° stimulus placed at a 6° eccentricity, with a 4 Hz temporal sinusoidal modulation. Closed circles indicate measurements made with the luminance of the background set throughout the test session. Very similar measurements were obtained (open circles) when the background luminance was set to equiluminant during the response period between stimuli presentations and only decreased (to black) or increased (to white) to coincide with stimulus onset. The similarity between the measurements obtained with these two different presentation parameters indicates that the effect of luminance in the surround is not due to slow adaptive mechanisms and that the surround provides fast feedback to the center.

### Supplementary figure 3

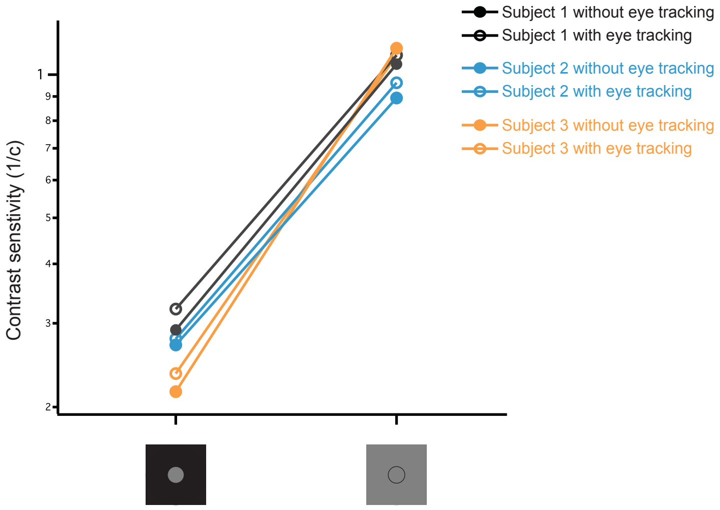


**Comparable contrast threshold measurements obtained with and without eye tracking.**
Temporal contrast sensitivity in three normal vision subjects each indicated with a different color. Subjects were asked to fixate a spot placed in the center of the CRT monitor. Stimuli were 1 degree diameter, 4 Hz sinusoidal temporal modulation presented at 6° eccentricity on either side of the fixation spot. Subjects were asked to indicate on which side the stimulus was presented. Their ability to detect the stimulus was determined by their forced choice response. Contrast sensitivity was reduced by a low luminance (0.15 cd/$m^2$) vs luminance matched (27.7 cd/$m^2$) surround. Closed circles indicate threshold measurements made without the aid of a pupil-tracking device. Very similar threshold values were obtained (open circles) when measurements were made with the aid of a pupil tracking device that automatically discounted trials if saccades were made during the presentation of the stimulus (see Methods).

### Supplementary figure 4

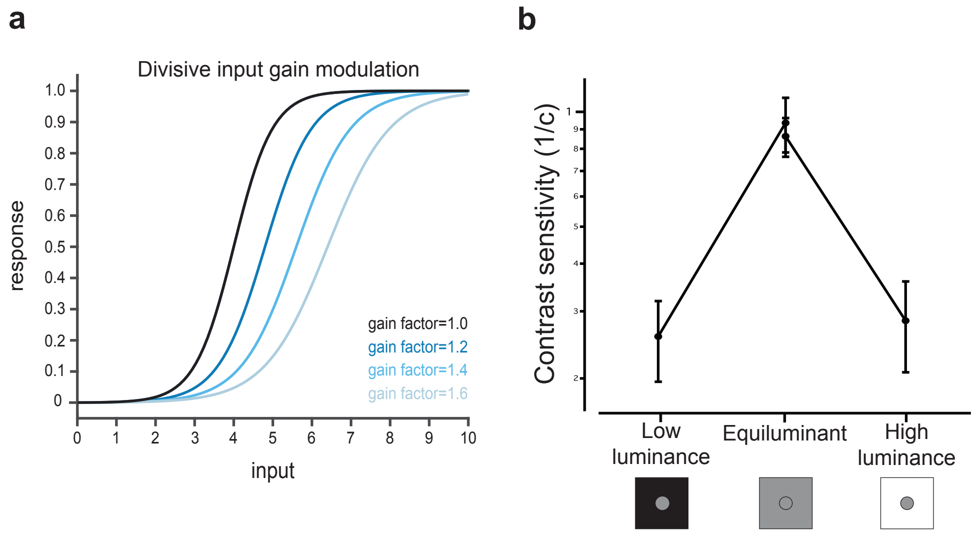


**Surround luminance acts as an adaptive mechanism**

**a**, Schematic showing input-output functions at different gain values (indicated by corresponding color), when the gain is implemented through a division of the input (compare this schematic to the results shown in Fig.1b). A divisive input gain mechanism scales and shifts the function along the input axis without scaling the function along the response axis, maintaining the proportionality between the slope of the function and the mean input value. **b**, A high luminance (64 cd/m$^2$) surround reduced contrast sensitivity similarly to a low luminance surround (n=9, error bars show standard deviation around a mean value).

### Supplementary figure 5

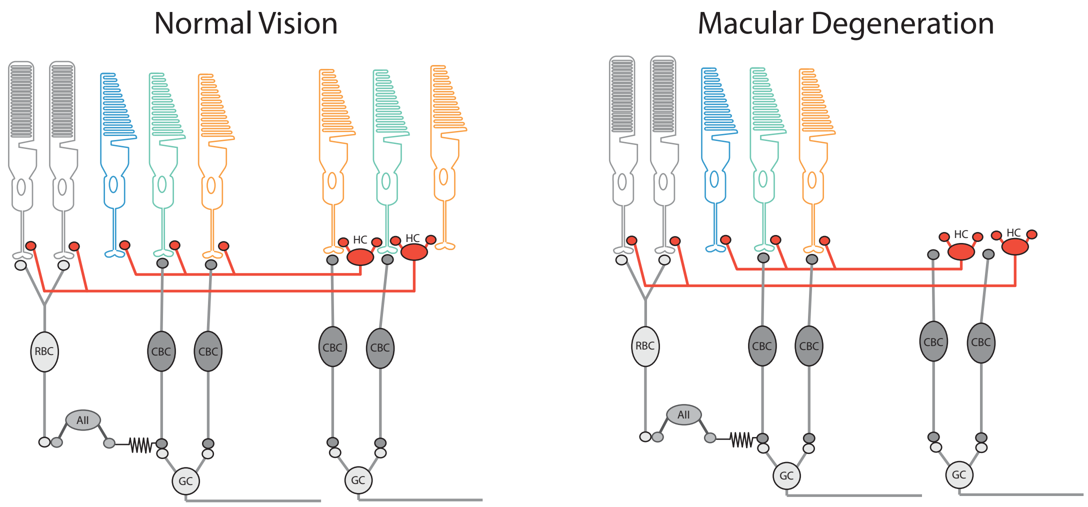


**Schematic of human retinal circuits showing horizontal cell feedback.** Horizontal cell (HC) feedback circuits are shown in red. In normal vision (left panel) HCs recieve input from L and M-cone photoreceptors and provide lateral feedback to all photorecptors. In conditions where there is local degeneration of photoreceptors for example in macular degeneration, cone input to horizontal cells and consequently lateral inhibition is missing. If this circuit had a suppressive function, re-activating lateral inhibition would further reduce vision in neighboring areas. Instead, the adaptive nature of the lateral inhibition means that reinstating this lateral feedback will improve vision in neighboring areas. S- and M- and L-cones are shown in blue, green and orange, respectively. Rod photoreceptors shown in grey. CBC: cone bipolar cell, RBC: rod bipolar cell, AII: AII amacrine cell, GC: ganglion cell.

### Supplementary figure 6

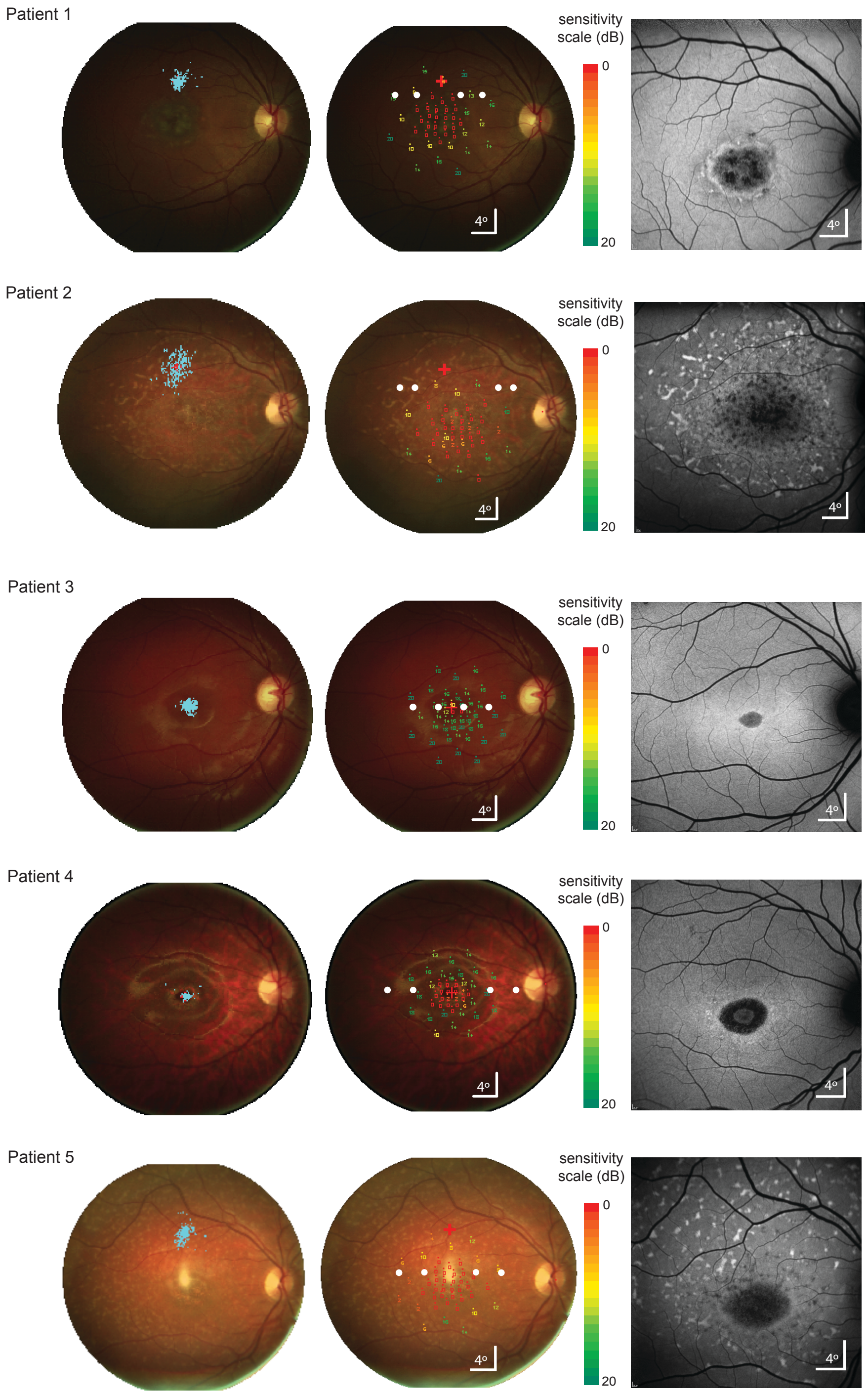


**Microperimetry data and Fundus Autofluorescence images for all tested Stargardt disease patients**.
Left column: fixation stability (in cyan, see Methods) superimposed on a color fundus image of the eye. Middle column: sensitivity values obtained from microperimetry superimposed on color fundus photograph of the eye. Color bar indicates sensitivity in decibels. Red cross indicates average fixation spot for images in left and middle column. Superimposed white circles indicate stimulus size and locations used for psychophysical testing of contrast sensitivity proximal (central placement) and distal (peripheral placement) to the lesion. Right column: short wavelength (486 nm) Fundus Autofluorescence images. The area of atrophy presents with a decreased signal in all patients.

### Supplementary figure 7

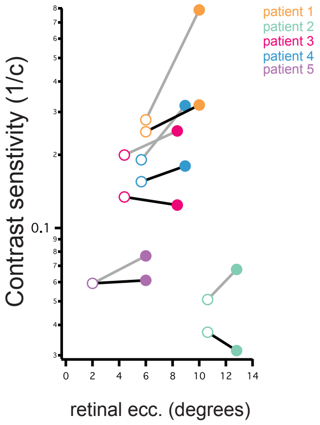


**Comparison of contrast threshold measuements in patients with Stargardt disease**.
Data for each patient is indicated with a different color. Measurements were made at locations close to the lesion and at a location further away from the sion lesion (open and filled circles, respectively) and when surrounded with low or equiluminant surrounds (measurements joined by black or grey lines respectively).

### Supplementary figure 8

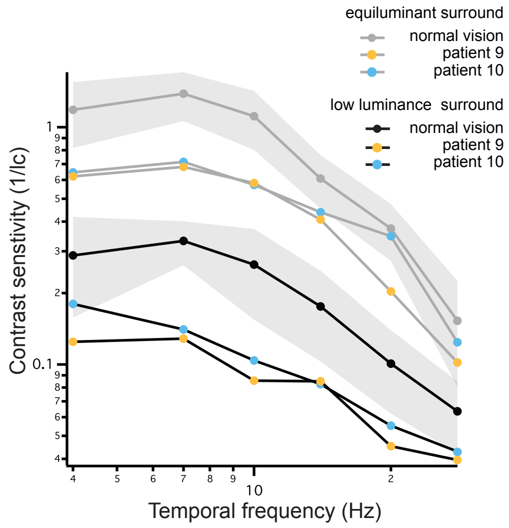


**Temporal contrast sensitivity functions for two patients with Congenital Stationary Night Blindness**
For both patients, contrast sensitivity was improved by an equiluminant (grey traces) vs low luminance (black traces) surround. Comparative data for normal vision subjects is shown (grey and black markers, n=8). Values shown are mean averages, standard deviation indicated by grey shading, data as shown in Fig. 1b).

### Supplementary figure 9

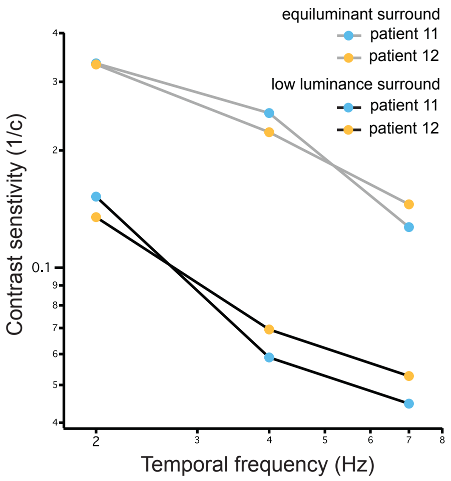


**Rod-only temporal contrast sensitivity functions for two patients with Bornholm Eye Disease.**
Silent substitution allowed selective stimulation of rod photoreceptors. For both patients, contrast sensitivity was worsened by a low luminance (black trace) vs equiluminant (grey trace) surround at all tested frequencies. (Data at 4hz also shown in Fig.1j,k)

### Supplementary figure 10

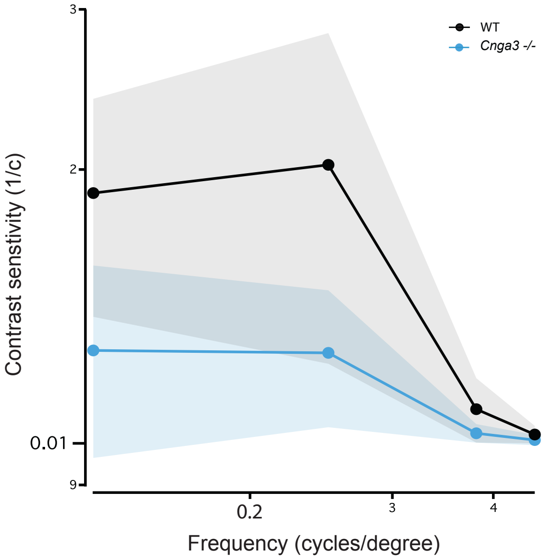


**Contrast sensitivity functions measured with a 1° height sinusoidal stimulus.** Contrast sensitivity functions could be obtained for wild-type (n=4, black trace) and *Cnga3*$^{-/-}$ mice (n=4, blue trace) using optomotor behavior driven by a 1° height sinusoidal stimulus. As expected, contrast sensitivity values were reduced compared to those obtained with full screen presentation of the same stimuli (Fig.3b). Values shown are mean averages, shaded regions denote standard deviations.

### Materials and Methods

#### Psychophysics experiments in human subjects

This study adhered to the tenets of the Declaration of Helsinki and was approved by the Moorfields Eye Hospital Ethics Committee (NHS REC reference: 11/H0703/10). Informed consent was obtained from subjects or consent and assent were obtained from parents and children, respectively, prior to entering the study. All psychophysical tests involved a 2-alternative forced choice task to detect the presence of stimuli on the left or right of a central fixation point. A chin/head rest maintained head position and viewing distance during the tasks. For a subset of subjects, comparative measurements were made with and without the aid of an eye tracking device (Tobii Pro X120). For these experiments, a calibration procedure was performed at the start of the session for each subject. The gaze outlier criterion was set at 100 pixels. The similarity between measurements made with and without pupil tracking suggested that pupil tracking did not significantly alter contrast threshold measurements and was therefore not used to generate the data presented in this study. This comparative data is presented in Suppl.Fig.3. Stimuli were presented via a gamma corrected CRT monitor, placed 0.8m from the chin rest. All tasks were performed binocularly, except for experiments in Stargardt patients for which the left eye was patched. Stimuli were generated in MATLAB (MathWorks inc, Natick, MA, USA) and delivered to the CRT monitor via a Visage MKII Stimulus Generator (Cambridge Research Systems Ltd, Rochester, UK). Contrast sensitivity thresholds were determined using a QUEST algorithm procedure (Watson and Pelli, 1983). All tests used a fixed number of reversals (minimum of 14 and a maximum of 28) to estimate a subject's threshold. Stimuli were presented for two seconds and subjects were allowed to respond without a timeout limit. Throughout the manuscript threshold values have been presented as contrast sensitivity (calculated as the inverse of the estimated threshold value). Normal vision subjects included 12 males and 10 females, aged 18-52. Three subjects (authors MR, KP and AG) completed all tests. All other subjects completed a subset of tests. Subjects had to fixate a fixation spot placed in the center of the monitor. Stimuli were presented at the same elevation as the fixation spot and at the specified and identical left or right eccentricity. Presentation either left or right was randomized and subjects were asked to report where they perceived the stimulus by pressing a left or right button on a console. For Stargardt patients the location of the stimulus presentation was chosen on the basis of microperimetry data (see below). For the majority of experiments the silent substitution technique (Estevez et al. 1982) was used to exclude S-cone activation. The rationale for this method was to negate possible differences arising from previously reported anatomical isolation of the S-cone pathway from interaction with rod input. All tests in patients with disrupted cone function (achromatopsia and blue cone monochromacy) were performed with non-selective stimuli modulated along a black-to-white axis. Comparative tests using black-to-white stimuli in normal vision subjects confirmed the main finding. Temporally modulated stimuli were presented as Michelson contrast sinusoids. A 0.2s ramp was used at the beginning of the stimulus. Gabor patches were used for spatially modulated stimuli. The range of contrasts presented varied between 0.1 and 30%. No measured threshold reached these minimum or maximum values. Patients with Bornholm eye disease only express three classes of opsin in the retina (in two of the patients tested, the expression was S-/M-/Rhodopsin and in the third, S-/L-/Rhodopsin). As our CRT monitor enables isolation of up to three classes of opsin we were able to generate rod photoreceptor-isolating stimuli for these patients, allowing us to specifically activate rods with our test stimulus. Surround annuli that contained more than one luminance were calibrated in order to have the same area dedicated to each luminance. For all annuli, 'width' was defined as the difference between outer and inner radius of the annulus.

A typical testing session with a single subject would normally allow measurement of 6-7 contrast sensitivity thresholds. No subject was tested for more than 1 hour in a single session. If a session consisted of testing a range of experimental values (for example a range of surround luminance values or stimulus frequency values) these were presented to the subject in pseudo-random order. Temporal contrast thresholds were measured using sinusoidally modulated stimuli at a range of cycle frequencies to build contrast sensitivity functions. Single measurement tests were performed at 4Hz cycle frequency. Spatial contrast thresholds were measured by presenting a Gabor patch with a range of fixed spatial frequencies, at varying contrasts. Conversely, spatial acuity measurements were performed by presenting a 40% contrast Gabor patch at varying spatial frequencies. In a small set of experiments, the surround changed only during presentation of the stimulus in the center (in order to test for/avoid adaptation to the mismatched surround).

In this case, the surround was equiluminant until the stimulus was presented and returned to equiluminant once the choice was made and the trial ended. Data from two subjects in which measurements were made with these presentation parameters is presented in Suppl. Fig.2.

**Microperimetry and Fundus Autofluorescence imaging in Stargardt disease patients**

Microperimetry was performed monocularly using the Nidek MP-1 (Nidek Technologies, Padova, Italy). Pupils were dilated and cyclopleged using 2.5% phenylephrine hydrochloride solution and 1% tropicamide ophthalmic solution. Before testing, the Spectralis OCT (Heidelberg Engineering, Heidelberg, Germany) was used to obtain a single transfoveal horizontal line scan. This was imported and used by the Nidek MP-1 manufacturer's software as an aid to automatically locate the anatomic fovea to facilitate accurate foveal placement of the testing grid. Testing consisted of a 4 apostilbs (1.27 cd/m2) background, Goldmann size III stimulus presented for a duration of 200 ms, and a 4 to 2 dB full-threshold bracketing test strategy. The customized testing grid consisted of 44 testing locations as previously described (Tanna et al 2018). The grid pattern was of radial design with centrally-condensed spacing and covered the macular and paramacular regions. All tests were performed under almost dark (mesopic) light conditions. The sensitivity at each retinal location was determined by iteratively adjusting the light intensity until the dimmest visible stimulus was found. The sensitivity for each test location was determined on a scale of 0 dB to 20 dB (MP-1 scale), with higher values indicating greater sensitivity. Only right eye data was used. Results from the microperimetry were used to estimate the functional border of the scotoma for each Stargardt patient. Test locations with 0-1 dB scores (ie, retinal locations where only the brightest stimulus was detected or no stimulus at all was detected) were defined as scotoma and stimuli were placed laterally adjacent to these locations for psychophysical testing. Before commencing the QUEST estimation of contrast sensitivity, we confirmed the suitability of the location by empirically testing the stimuli locations; patients were asked if they could see a 1 degree test stimulus (a static grey circle on a black background). If necessary, the test stimulus was moved laterally until the patient reported being able to clearly see it (location as shown in the figures). Fixation stability was determined prior to commencing microperimetry testing by sampling at 25 Hz for 5-15 s. Fundus images were acquired by white light reflectance imaging. Short wavelength (486 nm) fundus autofluorescence was obtained using a Heidelberg Spectralis Scanning Light Ophthalmoscope.

**Experiments in Mice**

All experiments in mouse were approved by the local Institutional Animal Care and Use Committees (UCL, London, UK) and conformed to the guidelines on the care and use of animals adopted by the Society for Neuroscience and the Association for Research in Vision and Ophthalmology (Rockville, MD, USA). C57/B6 mice were obtained from Harlan, UK. *Cnga3*$^{cpfl5/cpfl5}$ mice (referred to in the manuscript as *Cnga3*$^{-/-}$) were obtained from J.R. Heckenlively (University of Michigan).

**AAV vector production and testing**

A 3.0 Kb sequence from the mouse Gja10 promoter was amplified by PCR from mouse genomic DNA and cloned into an AAV viral vector containing the gene for Cre recombinase. AAV2/8 particles were produced and titered as previously described (Nishiguchi et al., 2015). Subretinal injections were performed in adult wild-type and *Cnga3*$^{-/-}$ mice. Tests for vector specificity were performed by subretinal injection in Ai9 (lox-STOP-lox-tdTomato, Allen Institute) mice. Two injections were performed in each eye (in the superior and inferior hemisphere respectively). 2μl of virus (titer $1x10^{13}$ vg/ml) were injected. When two viruses (pAAV-Gja10-Cre and pAAV-lox-STOP-lox-Hm4D) were co-injected, equal volumes of $2x10^{13}$ vg/ml were mixed in a vial prior to injection. Virus was diluted following purification and to the appropriate titer in sterile PBS-MK. Control injections were performed with the same volume of PBS-MK. For each mouse, one eye was randomly assigned virus injection and the other sham injection. All mice were allowed to recover for at least 3 weeks following subretinal injections before behavioural testing. For imaging of sparsely labeled horizontal cells, we made use of an existing dataset, where horizontal cells were labeled by subretinal injection of AAV2/2 vector in young adult wild-type mice. The AAV2/2-YC3.6. YC3.6 vector combines YFP and CFP and is a calcium sensor (Nagai T. et al., 2004), but it was used only for labeling purposes in this study. Quantification of horizontal cell transduction was performed by co-staining with a Calbindin antibody (Swant, dilution 1:500).

**Mouse optomotor behavior**

Mice were tested for Optomotor reflex in a setup (Cerebral Dynamics, NY, USA) consisting of 4 computer screens, simulating a rotating drum. An experimenter blind to the content of the subretinal injection performed scoring of the behavior. A clockwise vs counterclockwise 2 Alternative Forced Choice was used. The threshold was determined once the test completed 7 reversals within 1% contrast. All acuity tests were performed at a fixed 100% contrast. Contrast sensitivity tests were performed at the indicated spatial frequencies. Mice were allowed to adapt to the set-up platform for 10 minutes on the first day, before testing begun. All tests were repeated four times for each condition for each mouse. In order to allow activation of the chemogenetic tool Hm4Di, mice were intraperitoneally injected with a Clozapine-N-oxide compound between 1 and 3 hours before testing took place. 24 hours were allowed to pass before the following test was performed, in order to allow termination of CNO-mediated activation of the exogenous receptor.

Optomotor tests with a thin stimulus were performed in the same setup (with the stimulus presented on all four screens). The stimulus was generated by a custom Matlab script and consisted of a left- or right-ward drifting sinusoidal, placed in the center of the screen. A QUEST Staircase procedure was used to determine the contrasts to be presented to the mouse. The surround (consisting of the entire screen except for the 1 degree height stimulus) was either black, grey (coinciding with the midpoint of the sinusoidal stimulus) or white. To avoid adaptation to a different light level, all screens were grey during inter-trial intervals and the ‘surround’ was only presented simultaneously with the drifting stimulus.

**Histological analysis**

Eyes for histological analysis were taken post-mortem. Retinas were extracted and fixed in 4% PFA overnight. A histological clearing procedure (adapted from Costantini et al., 2015) was performed on whole retinas to maximize imaging quality and aid verification of expression of the fluorescent protein in Ai9 mice and mice co-injected with pAAV-Gja10-Cre and pAAV-EF1A-DIO-hM-4Di-EGFP. Images were obtained with a Leica confocal Microscope (model).

**Electroretinogram (ERG)**

The animals were anesthetized with an intraperitoneal injection of a 0.007 ml/g mixture of medetomidine hydrochloride (1 mg/ml), ketamine (100 mg/ml), and water at a ratio of 5:3:42 before recording. Pupils were fully dilated using 1.0% tropicamide. Subdermal ground was inserted in the mouse left cheek. A drop of Viscotears 0.2% liquid gel (Dr. Robert Winzer Pharma/OPD Laboratories, Watford, UK) was placed between the electrode and the eye. Bandpass filter cutoff frequencies were 0.312 Hz and 1000 Hz. Flicker series were performed at 0.1 cd/m$^2$ light intensity as previously reported (Nishiguchi et al., 2015). The frequencies tested were 1, 3, 6, 9, 12, 15 Hz. Three repetitions were performed before and three after CNO injection. Mice were moved and re-positioned after every repeat. Tests before and after CNO injection were performed on different days, as the induction time (1-3 hours) would not be covered by the same injection of anaesthetic.

**Statistics**

Unless otherwise stated averaged data is represented as mean ± standard deviation. The statistical tests used for each experiment are stated in the corresponding figure legend.

**Data Availability**

The data that support the findings presented in this study are available, upon reasonable request, from the corresponding authors.
